# Predicting Human Protein Function with Multi-task Deep Neural Networks

**DOI:** 10.1101/256420

**Authors:** Rui Fa, Domenico Cozzetto, Cen Wan, David T. Jones

## Abstract

Machine learning methods for protein function prediction are urgently needed, especially now that a substantial fraction of known sequences remains unannotated despite the extensive use of functional assignments based on sequence similarity. One major bottleneck supervised learning faces in protein function prediction is the structured, multi-label nature of the problem, because biological roles are represented by lists of terms from hierarchically organised controlled vocabularies such as the Gene Ontology. In this work, we build on recent developments in the area of deep learning and investigate the usefulness of multi-task deep neural networks (MTDNN), which consist of upstream shared layers upon which are stacked in parallel as many independent modules (additional hidden layers with their own output units) as the number of output GO terms (the tasks). MTDNN learns individual tasks partially using shared representations and partially from task-specific characteristics. When no close homologues with experimentally validated functions can be identified, MTDNN gives more accurate predictions than baseline methods based on annotation frequencies in public databases or homology transfers. More importantly, the results show that MTDNN binary classification accuracy is higher than alternative machine learning-based methods that do not exploit commonalities and differences among prediction tasks. Interestingly, compared with a single-task predictor, the performance improvement is not linearly correlated with the number of tasks in MTDNN, but medium size models provide more improvement in our case. One of advantages of MTDNN is that given a set of features, there is no requirement for MTDNN to have a bootstrap feature selection procedure as what traditional machine learning algorithms do. Overall, the results indicate that the proposed MTDNN algorithm improves the performance of protein function prediction. On the other hand, there is still large room for deep learning techniques to further enhance prediction ability.

## Background

The biological roles of the vast majority of known amino acid sequences remain partly or completely unknown: the UniProtKB database [1] currently stores more than 60 million sequences, but UniProt-GOA [2] lists only about 600 thousand experimentally-supported functional annotations in the form of Gene Ontology (GO) terms [3]. This information is far from uniformly spread across protein sequences, so elucidating their molecular activities, their whereabouts, their biological partners, and the environmental conditions enabling them has been increasingly dependent on computational methods that mostly perform annotation transfers from sequence [4]. Because naive or iterative application of these methods can generate uncontrolled error propagation in databases [5], curators nowadays rely on complementing transfers from orthologous proteins and from domain family assignments with mappings between controlled vocabularies and GO [2]. Notwithstanding, a substantial fraction of deposited sequences still has no annotations at all, many more lack information for at least one GO domain, and the hypotheses generated are often too generic to suggest a limited number of specific validation assays. Machine learning represents an attractive avenue to help fill in this gap, by modelling the relationship between protein function and the features extracted from individual or multiple biological data sources. When informative patterns can be detected, this approach can overcome the limitations of homology-based transfers due to the lack of similar sequences with known function, or to misleading alignment results. Many research groups have tested this hypothesis with success by examining heterogeneous data sources such as protein sequences [6–9], genomic information [10,11], gene expression profiles [12], and functional association networks [13], using neural networks (NN) [14], support vector machines (SVM) [7–9] and random forests [15]. Further efforts are going into integrative approaches, that try to leverage the strengths of individual methods and data types and to lessen the effects of their intrinsic limitations [16–19]. For instance, protein sequence and structure analysis can predict molecular function GO terms much better than biological process terms; in turn, the latter are more confidently inferred from genome-wide datasets. The evaluation results of the community-wide Critical Assessment of Function Annotation (CAFA) experiments confirmed these observations, but also highlighted that predicting protein function accurately still remains an open problem [20,21].

One of the major challenges supervised learning methods face in protein function prediction is the structured and multi-label nature of the problem, because the biological roles are described by sets of terms from the hierarchically organised GO domains. Given the complexity of this challenge and the tools at hand, most previous studies benefitted from handling many more tractable binary classification tasks [7–9] - one for each GO term. So far, very few groups have tried to build one classifier able to predict all relevant labels at once [14], but recent developments in the field of machine learning now make this approach feasible. Deep learning is a fast-evolving area of research, which tries to address regression or classification problems by extracting informative internal representations of the input data (aka feature representations) at different levels of abstraction. This is usually achieved with artificial deep neural networks (DNN), which include multiple hidden layers with hundreds of units each aimed at capturing high-level feature representations. The concept of DNN appeared a few decades ago, but remained impractical until Hinton and colleagues showed that layer-wise pretraining techniques allow deep networks to learn better feature representations and leading to improved classification performance[22]. Its growing power and popularity depend on several theoretical and technical advances that speed up training and reduce the risk of overfitting. Rectified activation functions force the model to learn sparse representations and ease vanishing-gradient issues that typically affect networks with many layers and sigmoid activation functions [23]. The dropout regularization technique - which consists in randomly omitting a different subset of the model parameters during training - greatly helps perform model averaging and reduce the risk of overfitting to known observations [24]. Finally, batch-normalization transforms each neuron input values from a randomly chosen subset of the training data (aka mini-batch) so that their distributions do not change dramatically during training [25]. The increasing availability of general-purpose graphical processing units (GPUs) and application programming interfaces (APIs) have also made a substantial contribution to the widespread application of deep learning techniques within both academia and industry. Deep learning has been applied to a wide range of problems in sequence and-omics data analysis, biomedical imaging and biomedical signal processing [26–29]. Multi-task deep neural networks are a particular type of architectures which consist of initial shared layers followed by as many independent modules (made up of individual output neurons possibly downstream of additional hidden layers) as the number of target labels (aka tasks). This design is meant to exploit commonalities and differences among tasks through the combination of shared and task-specific representations, and thus has the potential to compensate for the limited number of observations available for some tasks. This modelling approach has been previously applied to virtual screening [30], toxicity prediction [31] and prediction of protein biophysical features including secondary structure, solvent accessibility, transmembrane segments and signal peptides [32].

In this work, we investigate the usefulness of multi-task DNN (MTDNN) to tackle protein function prediction, an area which is expected to benefit from learning the dependencies among functional classes. We build a two-stage MTDNN structure, in which a set of feedforward layers shared by all tasks are in the first stage and as many task-specific feedforward layers as the number of tasks are parallel stacked upon the shared layers. This structure leverages both the shared representations of all tasks and specific characteristics of individual tasks. The effectiveness of the proposed approach is gauged against a naive multi-label DNN (MLDNN) - a feedforward structure with as many output units as the number of GO terms and several shared hidden layers-as well as three baseline methods. The experimental results show that MTDNN achieve higher *F*_1_ scores than the other methods tested; interestingly, the performance improvement over a single-task predictor is not linearly correlated with the number of tasks in the MTDNN model: medium size models with 20-50 GO terms appear to be more effective in our case. Another advantage is that there is no requirement for MTDNN to have a bootstrap feature selection procedure when given a set of features, in contrast to many traditional machine learning methods. This flexibility could be easily exploited in the future by adding complementary input features describing protein-protein interactions, gene expression profiles, literature co-occurrence, etc.

## Methods

### Data Collection

The protein sequences and feature data, the GO term vocabulary and the functional annotations used in this study are identical to those used to develop FFPred3 [9]. Annotations for human proteins were obtained from the Gene Ontology Annotation (GOA) [2] database released on 2015-02-02. The Gene Ontology (GO) OBO file released on 2015-02-03 was used for term definitions and semantic relations [33]. All GO terms in the biological process (BP), molecular function (MF) or cellular component (CC) domain with at least 150 annotations and 500 putative negative examples were selected. The negative examples of one GO term are the proteins that are (i) not labelled with the term under consideration, its descendants and its ancestors; (ii) nonetheless bear at least 2 MF terms and 2 BP terms with evidence code other than IC, NAS, TAS, and IEA. In total, 868 GO terms across the three domains were selected. Amino acid sequences were retrieved from UniProtKB version 2015_03 [1], and encoded through 258 features covering 14 different functional and structural aspects, including protein secondary structure, intrinsically disordered regions, transmembrane segments, signal peptides, post-translational modification sites, coiled-coil regions, and other sequence motifs [9]. These biophysical attributes can be easily calculated or predicted from amino acid sequences or their evolutionary profiles as reported before [8].

For benchmarking purposes, the set of human proteins that received GO term assignments supported by evidence code EXP, IDA, IMP, IGI, IEP, TAS or IC exclusively between 2015-02 and 2017-02 was collated from GOA database. Annotations to the term “*protein binding*” (GO:0005515) were discarded because they convey limited functional information. This test set was made up of 1754 annotations for 707 proteins in total – 349 MF annotations for 236 proteins, 556 BP annotations for 259 proteins, and 849 CC annotations for 492 proteins. Considering backpropagation, we have 2196 MF annotations covering 527 GO terms, 7353 BP annotations covering 1712 GO terms, and 4906 CC annotations covering 256 GO terms.

### Multi-task Deep Neural Networks (MTDNN)

#### Overview

MTDNN implements a multi-task architecture, with a set of feedforward layers shared by all tasks, upon which as many task-specific feedforward subnets as GO terms under investigation are parallel stacked - see Figure 1(a). This layout is meant to help the network learn individual tasks partially using a shared representation and the rest from task-specific characteristics. The network architecture is implemented using *Lasagne* and *Theano* [34]; each hidden layer is fully connected to the previous one, has batch normalised input values and has dropout applied in the course of training. The neurons in the hidden layers are activated by rectified linear functions, while output units make use of softmax functions with two outputs. The confidence score of each GO term corresponds to the value associated with the positive class from the relevant neuron.

**Figure 1.**
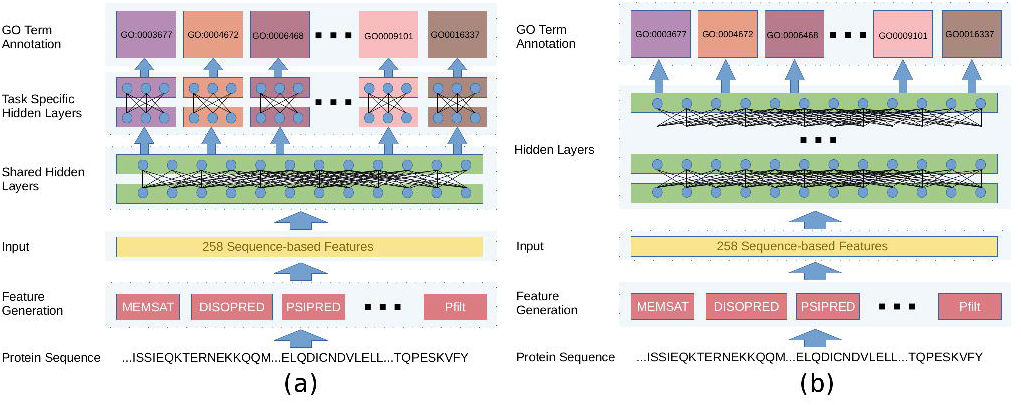
Schematic diagrams of MTDNN (a), and MLDNN (b).

#### Branching

We experienced difficulties in training one MTDNN network for each GO domain, because the high demands for memory exceeded the amount available on the GPUs. Therefore, we grouped the GO terms based on the “*is_a*” relationships in GO, and trained one separate model for each *branch* - one of the subgraphs rooted at each level-1 nodes - to predict the subset of its descendants included in our vocabulary. For example, our vocabulary lists 17 descendants of the level-1 term *immune system process* (GO:0002376), and the corresponding branch has 18 tasks, inclusive of itself. This procedure led to 38 branches in total: 18 within the biological process (BP) domain, 11 for molecular function (MF), and 9 for cellular component (CC). The summary of all branches is provided in supplementary material ST1. The number of GO terms in each branch varies a lot branch by branch. The largest branch is *biological regulation* (GO:0065007) with 218 output units for as many GO terms, while the smallest ones have only 2 output nodes. Given the semantic relationships in GO, some terms were included in different branches: *cell morphogenesis involved in neuron differentiation* (GO:0048667) appears in four branches, namely *cellular process* (GO:0009987), *single-organism process* (GO:0044699), *cellular component organization or biogenesis* (GO:0071840), *developmental process* (GO:0032502) for example. There are 290 out of 606 GO terms in BP, 71 out of 104 GO terms in CC, and 8 out of 158 GO terms in MF appearing in multiple branches, which generate as many predictions. For such GO terms, MTDNN returns one combined result by calculating the average value from the different branches.

#### Training and Optimisation

All tasks in MTDNN were trained individually while the shared layers were updated for all tasks. To this end, the protein sequences were initially clustered at 50% sequence identity with kClust [35], and for each task the resulting clusters were assigned to either the training or test set. Approximately 70-80% were used for training and the remaining 20-30% for testing - making sure there are at least 35 positive test examples for each GO term.

One of the most severe issues of protein function prediction using deep neural networks is the imbalance training problem, i.e., the numbers of positive examples in some GO terms are much fewer than the numbers of their negative examples. To deal with the issue, we investigated two strategies. The first one is that we aggregated weights of classes into the loss function to impose an additional cost on the model for making mistakes on the minority class during training. In this strategy, each batch composes of examples randomly sampled from the training set. The second strategy is that we oversampled the minority class, kept each batch with balanced positive and negative examples during the training, and in the prediction phase, penalized the minority class with a weight. The weight is defined as the ratio of the number of negatives to the number of positives in the training set. To determine the best strategy to deal with imbalanced training sets, we did a quick experiment only on MF terms because MF has a relatively small group of GO terms. The *F*_max_ performance of the strategy one is *0.244* and its *F*_1_ score at a threshold equal to 0.5 is 0.219, while the second strategy has obtained the *F*_max_ score *0.311* and *0.292* for its *F*_1_ score at threshold equal to 0.5. The results indicate that the strategy using balanced batch training and penalized inferring produced better results. Therefore, the results reported in the Results section were obtained using the balanced training strategy.

The training procedure consisted of 100 epochs and was divided into two stages: during the first 50 epochs both the shared and the specific layers were updated, while only the specific layers were modified during the last 50 epochs. This measure was aimed at reducing the degrees of freedom when task-specific layers were trained and at exploiting general feature representations of biological function to learn finer details. During each epoch, the GO terms were trained consecutively, and their order was randomly shuffled every time to minimise the risk of biasing the shared parameters towards more recently observed training examples.

Each MTDNN branch was independently optimised by searching a set of hyperparameters, including the number of shared layers, the number of hidden units in each shared layer, the number of specific layers, the number of hidden units in each shared layer, dropout rate, and learning rate - see supplementary material ST2 for more details. We employed the HYPEROPT package [36] to search the hyperparameter space randomly in 100 trials. The final models are taken from the parameters that maximize the average *F*_1_ score with threshold equal to 0.5 on the holdout test set.

### Multi-label Deep Neural Networks (MLDNN)

MLDNN implements a straightforward solution to multi-label problems and consists of a feedforward multilayer architecture with 258 input nodes (one for each sequence-derived feature), followed by several fully connected layers that are shared by one output layer - see Figure 1 (b). Each hidden layer has batch-normalized inputs combined through rectified linear units and is subject to dropout during training. The output layer is fully connected to the previous one, and consists of as many output neurons as the GO terms in the selected vocabulary activated by sigmoid functions, the output of which is returned as a confidence score. To train the MLDNN method, the protein sequences were first clustered at 50% sequence identity with kClust [35], and for each GO domain such clusters were assigned to the training or test set. Approximately 80% of the data were used for training and the remaining 20% for testing.

To optimise the MLDNN method, the set of hyperparameters in supplementary material ST3 was randomly sampled 100 times with the HYPEROPT package. For each trial, 100 epochs of training were carried out. The final models are taken from the parameters that maximize the average *F*_1_ score with threshold equal to 0.5 on the holdout test set.

### Single-task Deep Neural Networks (STDNN)

STDNN is a traditional divide-and-conquer solution of the multi-label learning problem, which employs a single fully connected feedforward deep neural network for an individual GO term, therefore, there are totally 868 binary DNNs optimised and trained independently. Like multi-task and multi-label implementations, the protein sequences were initially clustered at 50% sequence identity with kClust [35], and for each task the resulting clusters were assigned to either the training or test set. Approximately 70-80% were used for training and the remaining 20-30% for testing – depending on the number of positive annotations in each GO term and making sure there are at least 35 positive test examples for each GO term.

To optimise the STDNN, the set of hyperparameters in supplementary Table 3 was randomly sampled 100 times with the HYPEROPT package. For each trial, 100 epochs of training were carried out. The final models are taken from the parameters that maximize the average *F*_1_ score with threshold equal to 0.5 on the holdout test set.

### Baseline Methods

FFPred3 [9] was used to predict GO terms for the sequences in the benchmark set starting from the same 258 input features fed to both the MTDNN and MLDNN. Unlike the multi-task and multi-label deep learning approaches, however, FFPred3 examines the input values through a library of 868 Support Vector Machines independently trained to classify as many functional categories.

Naive predictions were generated based on the frequency of the GO term annotations for human sequences in UniProt-GOA released on 2015-02-02. The initial counts were obtained for all GO terms supported by the evidence codes EXP, IDA, IPI, IMP, IGI, IEP, IC and TAS. The data were then propagated following “*is a*” links in the GO released on 2015-02-03, and scaled between 0 and 1 for each domain separately, by dividing the final counts by the number of occurrences of the root node.

BLAST predictions were obtained by collecting all BLAST hits in the UniRef90 sequence database released on 2015-02 with an E-value greater than 1e-03. Then the annotations in UniProtKB released on 2015-02-03 supported by evidence codes EXP, IDA, IPI, IMP, IGI, IEP, IC and TAS were transferred to the target sequences. The confidence scores of GO terms were calculated by dividing the local alignment sequence identity by 100. When multiple BLAST hits were annotated with the same GO term, the highest score was retained.

### Performance Evaluation

Prediction accuracy was measured by protein-centric precision-recall analysis separately for each GO domain. For each protein ***x*** in the benchmark set and decision threshold *t*, the set of predicted GO terms ***G***_***x*, *t***_ was built by collecting all terms with confidence scores greater than or equal to *t*, and their ancestors in GO linked by “*is_a*” relationships and different from the root. The precision *p*_*x*, *t*_ and recall *r*_*x*, *t*_ can be respectively written as

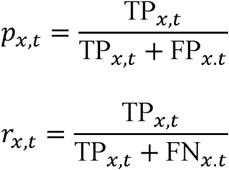

where TP_*x*, *t*_ is the number of true positives, FP_*x*, *t*_ is the number of false positives, and FN_*x*, *t*_ is the number of false negatives for the benchmark protein *x* at threshold *t*. Then the average across the test set are taken as

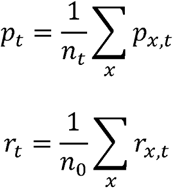

where *n*_*t*_ is the number of target proteins with at least one prediction scoring above threshold *t*, and *n*_*0*_ is the number of target proteins in the GO domain in the benchmark set. Therefore, the average *F*_1_ for the threshold *t* and *F*_max_ were calculated as

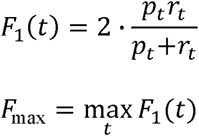

We also employed term-centric evaluation to measure *F*_1_ scores for individual GO terms covered by the benchmark set.

## Results and Discussions

We are interested in knowing whether, how and why multitask deep neural networks improve the performance of protein function prediction. In this section, we will show both holdout and benchmark evaluation results of our optimised MTDNN models, and discuss the lesion we learned from our experiments.

### Optimised Models

As detailed in Methods section, the hyperparameter optimisation procedure was carried out by using the Python HYPEROPT package, which randomly samples from pre-defined hyperparameter space. Each branch has a set of optimised hyperparameters including the network architectures like the depths of shared layers and specific layers, the numbers of hidden units in each layer *etc*., and the other non-architecture hyperparameters like the learning rates and regularisation options. The question that interests us most is what network architecture each branch chose in the optimisation procedure. We show the depths of both the shared layers and the specific layers for all branches of three domains as stacked bar charts in Figure 2. The branches are presented in a descending order from the top to the bottom. The red number on the right-hand side of each bar is the number of GO terms in the branch. We notice that the larger branch tends to choose deeper models in general, however, we are unable to observe any correlation between the depths and the branch sizes.

**Figure 2.**
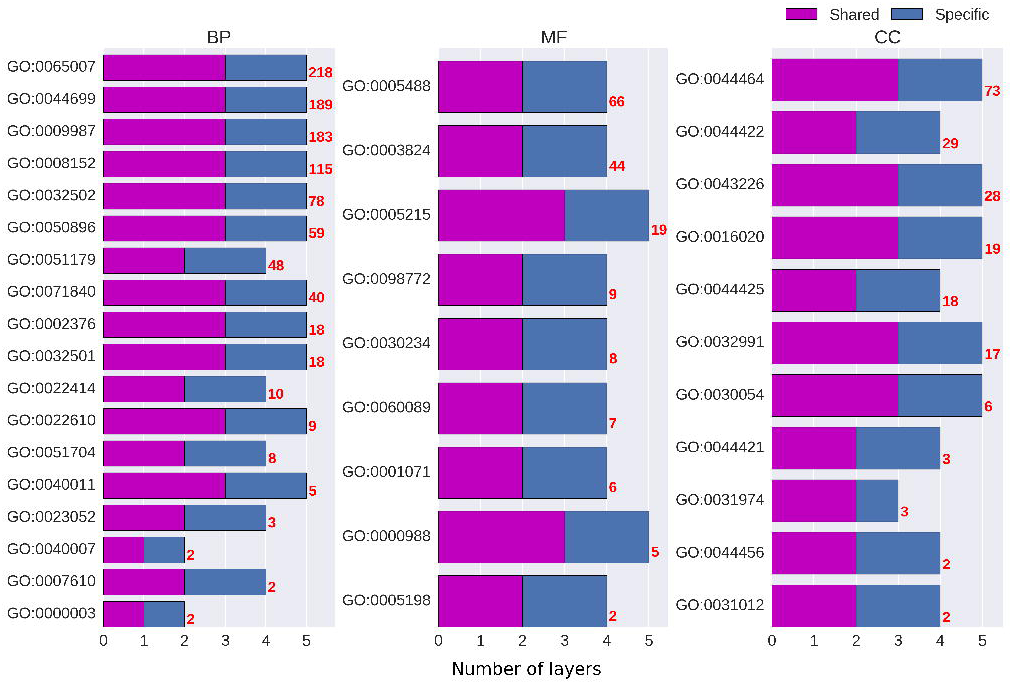
The numbers of layers of the optimised MTDNN models in BP, MF and CC domains. The red numbers on the right-hand side of bars represent the numbers of GO terms in individual branches.

We also show the summary of the numbers of hidden units in each layer of all optimised models as pie charts in Figure 3. Less than half branches chose three shared layers and 5% models have only one shared hidden layer. 47% of the models chose 800 hidden units in their first shared layer and 79% chose 500 hidden units in their second shared layer. 79% of the models chose 500 hidden units in their first specific layers and the same percentage of models chose 300 hidden units in their second specific layers. This result tells us that larger and deeper models are favourable in many branches according to our experiments.

**Figure 3.**
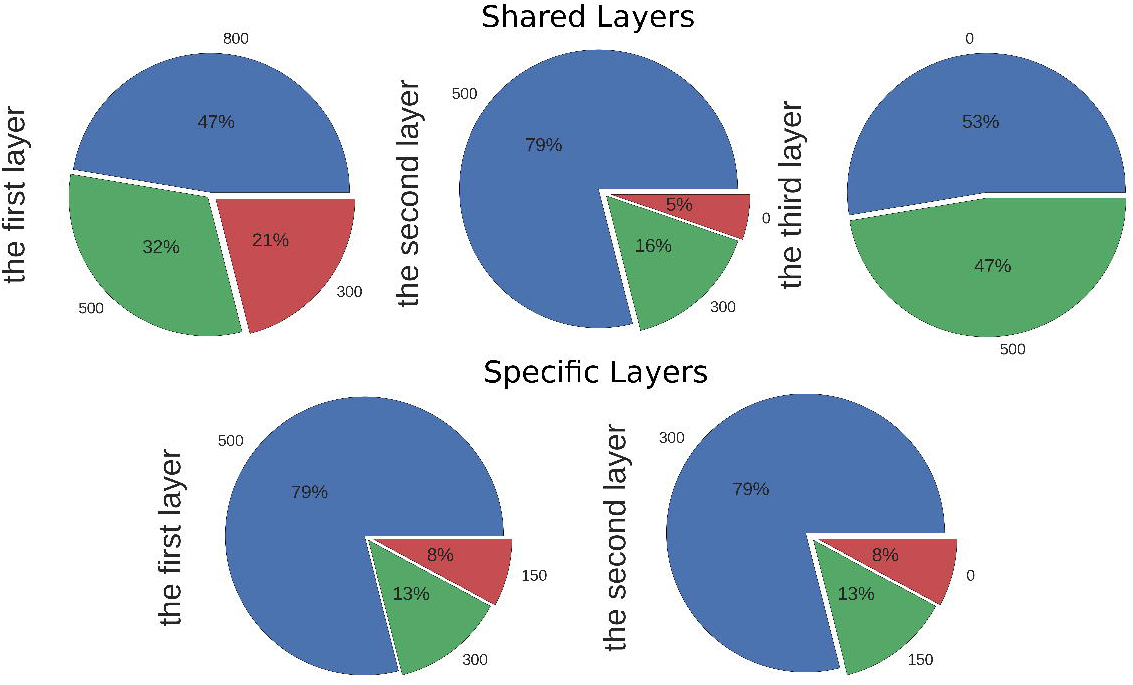
The summary of the numbers of hidden units in individual shared and specific layers of the optimised MTDNN models. The upper row presents the choices of numbers of hidden units in the three shared layers; the lower row presents the choices of numbers of hidden units in the two specific layers

### Holdout Set Evaluation

We only compare the performance of MTDNN with FFPred in the holdout evaluation because the generation procedure of holdout sets for MLDNN is totally different compared to MTDNN and FFPred. We evaluated term-centric *F*_1_ scores for both MTDNN and FFPred and considered the average *F*_1_ score differences between MTDNN and FFPred. In particular, we define term-centric *F*_1_ score the difference as:

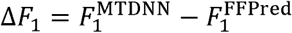

In Figure 4, we show the average *F*_1_ score differences for two groups of GO terms – all terms in our vocabulary and specific terms -- in BP, MF, and CC domains. The definition of specific terms in our case are those terms in our vocabulary which either have not child term at all or have children terms but they are not in our vocabulary. There are mainly two observations from this figure: firstly, generally speaking, MTDNN has better performance than FFPred, especially a considerable improvement in BP; secondly, though MTDNN overall improves the performance, specific terms improved less than general terms.

**Figure 4.**
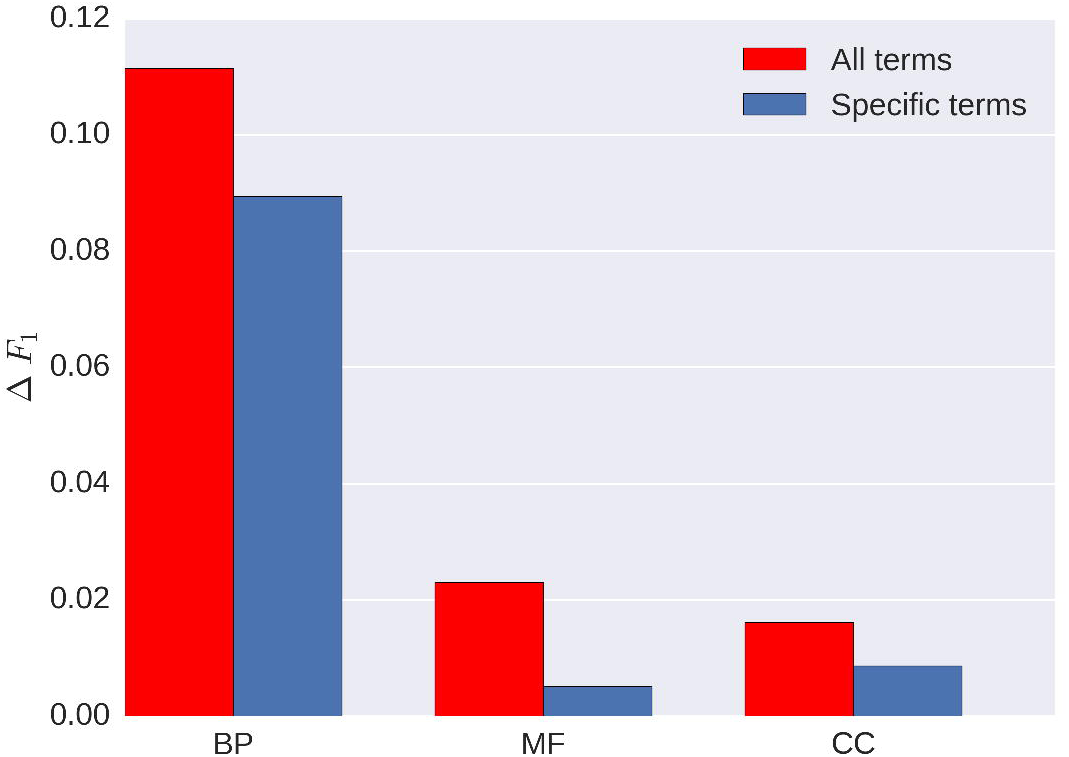
Average *F*_1_ score differences between MTDNN and FFPred in BP, MF and CC domains. The red bars represent the performance of all GO terms in our vocabulary and the blue bars represent the performance of specific terms.

Another question we should like to address is whether larger branches improve more than smaller branches, i.e. a model with more tasks gains more improvement from using MTDNN than a model with fewer tasks. This would be expected if MTDNN can effectively exploit both the shared and task-specific representations that are learnt from the different training sets. To address this question, we show the average *F*_1_ score differences for individual branches in Figure 5. The branches are ordered ascendingly from the left to the right in terms of their sizes and the red numbers on the top of bars are the numbers of GO terms in individual branches. In general, on one hand, small branches perform poorly, on the other hand, it does not seem to be true that the larger the branch is, the better it performs. There is no clear pattern that the performance improvement correlates with the size of the branch overall. In BP, there indeed is a weak pattern that branches with larger size tend to perform better, but there is no such pattern in MF and CC. We believe that it is a complex question combining other factors like the numbers of positive and negative examples in each GO term of each branch.

**Figure 5.**
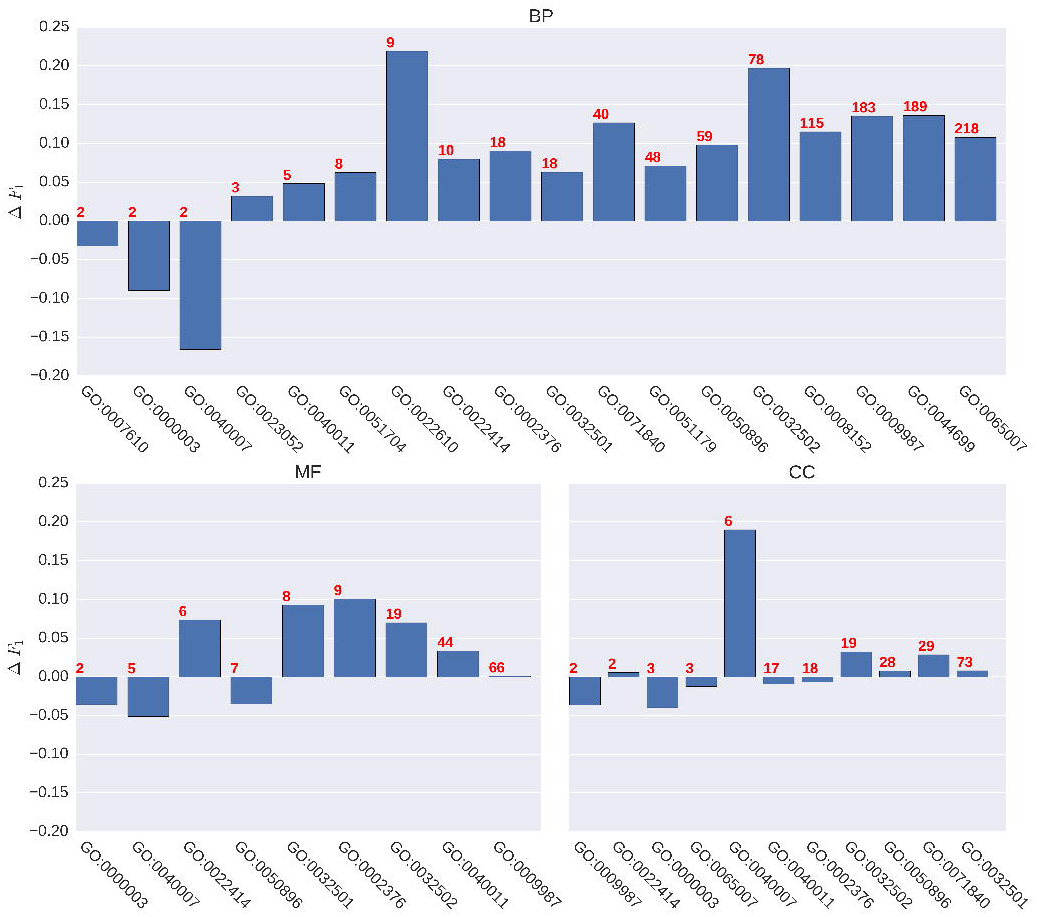
Average *F*_1_ score difference between MTDNN and FFPred for all branches in BP, MF and CC domains. The red numbers on the top of bars represent the numbers of GO terms in individual branches.

### Benchmark Set Evaluation

Here, one important question we need to address is, whether the MTDNN method is better than other methods in the benchmark set. To answer this question, we firstly evaluated the *F*_max_ performance of all compared methods following the practice in CAFA [20,21]. We report the results of five methods at the decision thresholds that maximize the *F*_1_ scores for each GO domain using the benchmark set in Table 1. The BLAST and naive methods are the two providing the worst *F*_max_ performance. Note that naive method has a good *F*_max_ performance in CC only because of the fact that the majority of proteins locate in the cytoplasm is biased towards the naive frequency counting method. More interestingly, the results indicate that the *F*_max_ performance of MTDNN is better than the other methods in BP and CC, but worse than FFPred, MLDNN and STDNN in MF. We notice that the thresholds at which other methods produce the *F*_max_ scores are farther away from 0.5 than MTDNN. Although *F*_max_ is commonly used to evaluate classification algorithms, it arguably is a paradox because in the reality the threshold to produce the maximum *F*_1_ score would never be known without knowing true labels of all predictions.

**Table 1.** *F*_max_ performance comparison between MTDNN and other prediction methods. For the five methods, the table reports *F*_max_ and the threshold values that produce the *F*_max_ in three GO domains, namely BP, MF, and CC. The best *F*_max_ scores in respective domains are marked in bold font.

| Methods | BP |  | MF |  | CC |  |
| --- | --- | --- | --- | --- | --- | --- |
| | *Threshold* | $F_{\max}$ | *Threshold* | $F_{\max}$ | *Threshold* | $F_{\max}$ |
| MTDNN | 0.52 | **0.298** | 0.58 | 0.311 | 0.54 | **0.484** |
| MLDNN | 0.26 | 0.287 | 0.30 | 0.343 | 0.23 | 0.449 |
| STDNN | 0.99 | 0.288 | 0.99 | 0.338 | 0.95 | 0.396 |
| FFPred | 0.80 | 0.293 | 0.85 | **0.355** | 0.86 | 0.477 |
| BLAST | 0.25 | 0.106 | 0.22 | 0.126 | 0.21 | 0.212 |
| Naive | 0.14 | 0.252 | 0.23 | 0.266 | 0.45 | 0.474 |

A more realistic evaluation is based on standard measure of binary classification accuracy – i.e. interpreting the scores as probability estimates and therefore setting the decision threshold equal to 0.5. The results under these evaluation settings in Table 2. Three metrics, namely the average precision, average recall, and average *F*_1_ scores are presented for the three GO domains. In terms of the *F*_1_ score, the best performing method out of five methods is MTDNN. Its *F*_1_ scores are better than other methods across all three domains. The results reveal that naive method and BLAST are still the two poorest methods. The results also show that FFPred and STDNN have the highest recall scores in all three domains, but with lower precision scores; on the contrary, MLDNN offers higher precision scores, but lower recall scores, in turn, lower *F*_1_ scores. This observation reveals that FFPred and STDNN made more predictions which on the one hand increases true positives, but on the other hand also increases false positives; while MLDNN makes fewer mistakes by reducing the number of predictions. MTDNN provides a balance between precision and recall keeping higher *F*_1_ in all three domains, i.e. MTDNN is able to reach a trade-off between predicting accurately and making more predictions.

**Table 2.** Performance comparison at threshold equal to 0.5 between MTDNN and other prediction methods. This table reports average *F*_1_, Precision, Recall scores of five methods in three GO domains, namely BP, MF, and CC, with the threshold equal to 0.5. MLDNN* is the performance obtained at the thresholds that maximise the *F*_1_ scores for individual GO terms in the training set. The best *F*_1_ scores in respective domains are marked in bold font.

| Methods | BP |  |  | MF |  |  | CC |  |  |
| --- | --- | --- | --- | --- | --- | --- | --- | --- | --- |
| | Precision | Recall | $F_1$ | Precision | Recall | $F_1$ | Precision | Recall | $F_1$ |
| MTDNN | 0.282 | 0.312 | **0.296** | 0.283 | 0.303 | **0.292** | 0.389 | 0.626 | **0.48** |
| MLDNN | 0.353 | 0.103 | 0.160 | 0.335 | 0.170 | 0.267 | 0.402 | 0.308 | 0.349 |
| STDNN | 0.070 | 0.676 | 0.127 | 0.112 | 0.606 | 0.189 | 0.168 | 0.858 | 0.281 |
| FFPred | 0.161 | 0.492 | 0.242 | 0.141 | 0.558 | 0.225 | 0.248 | 0.820 | 0.380 |
| BLAST | 0.129 | 0.019 | 0.033 | 0.447 | 0.035 | 0.066 | 0.423 | 0.027 | 0.051 |
| Naive | 0.539 | 0.024 | 0.046 | 0.318 | 0.117 | 0.171 | 0.446 | 0.407 | 0.425 |

Next, we used the benchmark annotations to assess the difference in prediction accuracy between MTDNN and FFPred, grouped the GO terms according to the size of the MTDNN branches in which they locate, and calculated the mean value of Δ*F*_1_ of GO terms in each branch. A graphical summary of this analysis is in Figure 6. Like the observations in holdout set evaluation, the performance improvement is not always linearly correlated with the number of tasks. Arguably, selecting the best choice of the size of the model is a data-dependent question and is not trivial. In our case, the range between 20 and 50 tasks in one model provides more improvement. However, breaking larger branches into smaller branches with 20-50 tasks, which sounds like a sensible solution, comes at the cost of reduced training sets, thus limiting the viability of deep learning approaches. Future work will research the best way into to develop a model using an expanded vocabulary and new datasets for our web service in the future.

**Figure 6.**
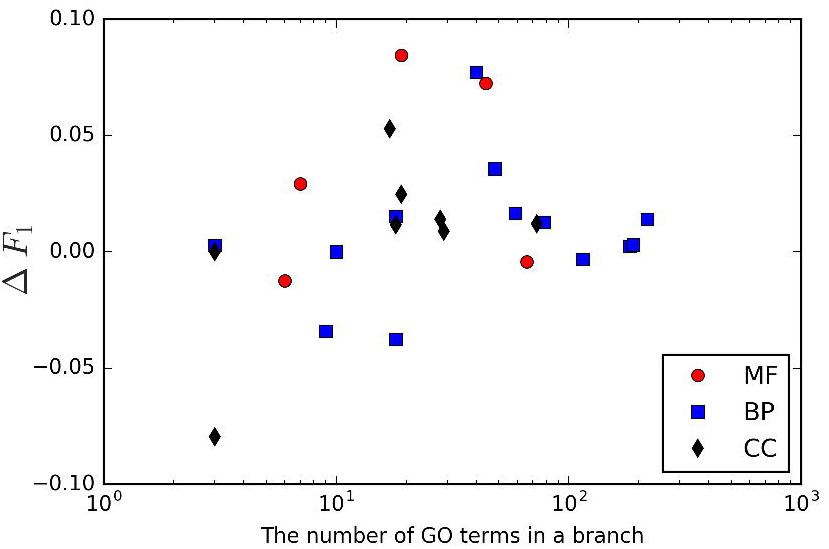
the scatter plot of mean values of *δF*_1_ against the number of GO terms in a branch.

## Conclusions

In this paper, we developed a multi-task deep neural network (MTDNN) architecture to tackle the multi-label problem in protein function prediction. MTDNN is able to learn both a shared feature representation from all GO terms and specific patterns from individual terms by employing two stacked multi-layer structures, one shared by all tasks and another one specific to each task on top of the shared one. Importantly, it is no requirement for MTDNN to proceed a bootstrap feature selection as what many traditional machine learning algorithms usually do. We compared MTDNN with five baseline methods, namely naive method, BLAST, FFPred, STDNN, and naive MLDNN. We then evaluated the accuracy of the proposed MTDNN using both holdout set and benchmark set. The results show that MTDNN offers better performance in holdout set evaluation, and also better performance on the benchmark set at the decision threshold equal to 0.5 by balancing precision and recall, which makes MTDNN more favourable in practical use than the other methods tested. Another interesting result is that the performance improvement of MTDNN over the single-task predictor is not always linearly correlated with the number of tasks in the model. In our case, medium size models provided more improvement. Encouraged by the success in the current study, which suggests that MTDNN is a better solution to tackle the multi-label problem, we intend to improve the performance of predicting protein functions further by adding complementary input features describing protein-protein interactions, gene expression profiles, or literature co-occurrence.

## Authors’ contributions

D.T.J. conceived the study and designed the experiments with R.F., D.C. and C.W.; R.F. conducted the experiments; R.F., D.C., and C.W. analyzed the data; R.F. and D.C. wrote the manuscript; all authors read and approved the final version.

## Acknowledgements

The authors are grateful to the members of the UCL Bioinformatics Group and to Jaqui Hodgkinson, Chris Cheadle, and Hongbao Cao from Elsevier for valuable discussions. This work made use of the high-performance computing facility of the Department of Computer Science at University College London. R.F. acknowledges funding from Elsevier; D.C and C.W. are supported by the UK Biotechnology and Biological Sciences Research Council (References: BB/L020505/1 and BB/L002817/1).

## Supporting information

**ST1. Summary of all branches**. This table reports all branch root terms, the number of GO terms in each branch, the names of branch root terms and their domains.

ST2. Hyperparameters and their value space for MTDNN. *Different layers may have different number of hidden units.

ST3. Hyperparameters and their value space for MLDNN.

